# Constitutive Inhibition of Transient Receptor Potential Canonical Type 6 (TRPC6) by O-GlcNAcylation at Threonine-221

**DOI:** 10.1101/2022.06.15.496344

**Authors:** Sumita Mishra, Junfeng Ma, Masayuki Sasaki, Federica Farinelli, Richard C. Page, Mark J. Ranek, Natasha Zachara, David A Kass

## Abstract

Transient receptor potential canonical type 6 (TRPC6) is a non-voltage gated cation channel that principally conducts calcium to regulate signaling in cardiac, vascular, neuronal and other cells. Abnormally increased TRPC6 expression/conductance and genetic gain of function mutations contribute to fibrosis, hypertrophy, proteinuria, and edema, notably linked to its stimulation of nuclear factor of activated T-cells (NFAT) signaling. Hyperglycemia (HG) also activates TRPC6/NFAT as a cause of diabetic renal disease. While prior work linked HG-TRPC6 activation to oxidant stress, the role of another major HG modification - O-GlcNAcylation, is unknown. Here we show TRPC6 is constitutively O-GlcNAcylated, TRPC6 and O-GlcNAc transferase proteins interact, this modification potently suppresses basal channel conductance and NFAT activity, and it is unaltered by HG. Proteomics identifies O-GlcNAcylation at Ser14, Thr70, and Thr221 in the N-terminus ankyrin-4 (AR4) and neighboring linker (LH1) domains of TRPC6. Of these, T221 is most impactful as a T221A mutation increases basal NFAT activity 11-fold, TRPC6 conductance 75-80% vs wild-type, and when expressed in cardiomyocytes amplifies NFAT-pro-hypertrophic gene expression. T221 is highly conserved and mutating homologs in TRPC3 and TRPC7 also markedly elevates basal NFAT activity. Molecular models predict electrostatic interactions between T221 O-GlcNAc and Ser199, Glu200, and Glu246, and we find similarly elevated NFAT activity from alanine substitutions at these coordinating sites as well. Thus, O-GlcNAcylation at T221 and its interaction with coordinating residues in AR4-LH1 is required for basal TRPC6 channel conductance and regulation of NFAT.

## INTRODUCTION

Transient receptor potential canonical type-6 (TRPC6) is a non-voltage gated transmembrane cation channel conducting primarily Ca^2+^ (Dietrich and Gudermann, 2014). It is widely expressed with generally low basal conductance, but its stimulation impacts podocyte function (Reiser et al., 2005), cardiac hypertrophy (Onohara et al., 2006; Seo et al., 2014b), myofibroblast differentiation (Davis et al., 2012), pulmonary vascular tone (Harraz and Jensen, 2021; Tauseef et al., 2012; Weissmann et al., 2006), wound healing (Davis *et al*., 2012; Takada et al., 2014) immune cell (Ramirez et al., 2018; Schilling and Eder, 2009) and platelet function (Paez Espinosa et al., 2019), and neuronal growth (Liu et al., 2020a). Selective inhibitors(Lin et al., 2019) are in human trials with interest in cardiovascular, renal, and pulmonary disease.

TRPC6 is primarily stimulated by GPCR signaling *via* diacylglycerol and phospholipase C (Dietrich and Gudermann, 2014), and by mechanical stretch (Seo et al., 2014a). Ca^2+^ influx via TRPC6 stimulates calcium-calmodulin activated phosphatase calcineurin (CaN) to dephosphorylate nuclear factor of activated T-cells (NFAT) triggering its nuclear translocation to alter gene expression (Kuwahara et al., 2006; Wang et al., 2015). NFAT consensus sequences in the TRPC6 promotor provides positive feedback to amplify its impact. Human TRPC6 gain of function mutations occur in familial segmental glomerulosclerosis (Hall et al., 2019; Santin et al., 2009; Winn et al., 2005b), with most mutations in the cytoplasmic N-terminus ankyrin repeat domains needed for channel assembly and activity (Azumaya et al., 2018; Tang et al., 2018).

TRPC6 conductance and associated NFAT activation is also controlled by post-translational modifications (PTM) (Hagmann et al., 2018; Hisatsune et al., 2004; Kanda et al., 2011; Liu et al., 2020b). For example, oxidant stress coupled to NADPH oxidases NOX2 and NOX4 stimulates TRPC6 signaling (Chaudhuri et al., 2017; Wang et al., 2009). N-glycosylation at N472 and N561 (Talbot et al., 2019) and phosphorylation at S14(Hagmann *et al*., 2018) are required for normal TRPC6 membrane localization and activity. Phosphorylation can both increase (T487 by calcium-calmodulin activated kinase, CaMKII) (Erickson et al., 2008), or decrease (T10, S262 by cGMP-activated kinase, cGK1α)(Kinoshita et al., 2010; Koitabashi et al., 2010) channel conductance and associated signaling.

A known stimulator of TRPC6 and associated NFAT signaling is hyperglycemia (HG), as models of diabetes exhibit greater TRPC6 expression, conductance, and associated NFAT activity in kidney (Ilatovskaya et al., 2018; Ma et al., 2019; Wang et al., 2010), monocytes (Ma et al., 2015; Wuensch et al., 2010), and platelets (Liu et al., 2008). This raises the possibility that the PTM mediated by O-linked β-N-acetyl glucosamine (O-GlcNAc) that increases by HG with diabetes (Ducheix et al., 2018; Fricovsky et al., 2012; Ma and Hart, 2013) and is linked to CAMKII activation, cardiac disease and arrhythmia (Erickson et al., 2013; Lu et al., 2020), also plays a role. This PTM is controlled by O-GlcNAc transferase (OGT) catalyzing addition of O-GlcNAc onto serine or threonine residues, and O-GlcNAcase (OGA) to remove it (Chatham et al., 2021; Hart et al., 2007), and it alters protein-protein interactions, structure, activity, subcellular localization, stability and degradation (Bond and Hanover, 2013; Chatham et al., 2020; Ma et al., 2022). Based on this, we speculated HG may stimulate TRPC6 in part by O-GlcNAcylation of the channel. We report quite the contrary, that TRPC6 is constitutively O-GlcNAcylated and without this basal channel conductance and NFAT signaling are markedly increased. The primary regulating residue is Thr221 in the 4^th^ ankyrin domain is, and its O-GlcNAcylation coordinates with residues at Ser199, Glu200, and Glu246 to constrain basal TRPC6 conductance and NFAT signaling in the physiological range.

## RESULTS

### NFAT activity is increased by hyperglycemia but not via enhanced O-GlcNAcylation

To explore the impact of HG on NFAT, TRPC6, and O-GLcNAcylation, we expressed promotor luciferase-reporter vectors in HEK-293T cells that intrinsically lack TRPC6, and determined the impact of HG. In non-transfected HEK cells, HG stimulated a modest rise in NFAT promotor activity, but this was significantly amplified by TRPC6 expression (**Figure 1A**) confirming engagement of TRPC6 in the response. Mannitol served as a negative control. HG also dose-dependently stimulated TRPC6 promotor activity (**Figure 1B**) consistent with its NFAT-activated promotor. HG broadly increased protein O-GlcNAcylation, and this was mimicked by incubating cells with either the OGA inhibitor Thiamet-G (TMG) or stimulation of UDP-GlcNAc synthesis with glucosamine (GlcN, **Figure 1C, 1D**). However, unlike HG, neither TMG nor GlcN increased NFAT activity above baseline (**Figure 1E**), indicating that O-GlcNAcylation did not activate TRPC6-dependent NFAT per se.

**Figure 1.**
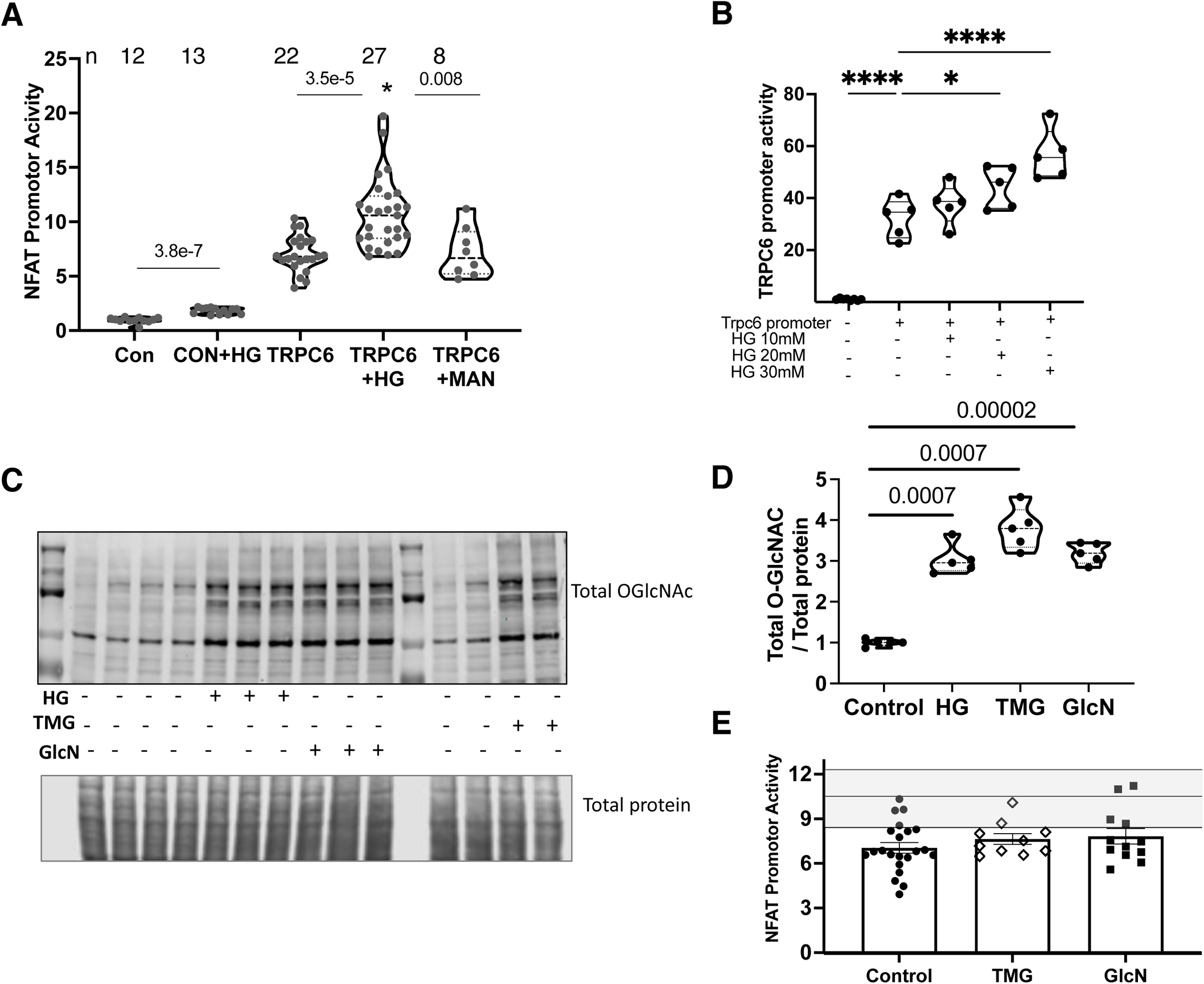
Influence of hyperglycemia (HG) on NFAT and TRPC6 promoter activity and protein O-GlcNAcylation. **A)** NFAT promotor activation by HG in HEK-293T cells lacking or expressing TRPC6. Mannitol (Man) serves as control. (n=12-27, P value for −TRPC6 (CON) +/− HG is Mann Whitney U test; P values for +TRPC6 +/− HG or Man are from Kruskal Wallis test with Dunns multiple comparisons test (MCT). *\** - P=0.01 for interaction of HG and TRPC6 expression by 2W-ANOVA. Sample size for each shown at top. **B)** Dose dependent TRPC6 promoter activity in response HG (1WANOVA, Dunnett’s MCT; *P =0.035, ****P < 0.0001, n = 8 for control, n=5 for other groups. **C)** Western blot for total O-GlcNAc in lysates from HEK-293T cells treated with 30 mM HG, +/− 10 μM TMG or 10 μM Glucosamine (GlcN) for 6h. **D)** Quantitation of data from experiment C. (n=5/group, Welch ANOVA, P-values Dunnet’s MCT) **E)** Increase in NFAT promoter activity in TRPC6 transfected cells treated with vehicle, 10 μM TMG or 10 μM glutamic acid (GlcN). Each response is significantly below than of TRPC6 expressing cells +30 mM HG (shaded area – median, 25, 75%, P<0.003 for each group comparison to HG response by 1WANOVA, Dunnett’s MCT).

### TRPC6 is constitutively O-GlcNAcylated in the N-terminus Cytoplasmic Domain

The preceding findings could be explained if O-GlcNAcylation did not impact TRPC6 modulated NFAT stimulation, or if TRPC6 was already modified in this way so that further increases had negligible effects. To test this, HEK-293T cells were transfected with TRPC6-YFP fusion to facilite expression detection and provide a robust epitope for immuno-precipitation (IP). IP was performed with either YFP or O-GlcNAc as bait, and the precipitate probed with anti-GFP (serves as anti-YFP) and monoclonal anti-O-GlcNAc antibodies. This revealed recombinant TRPC6 (~130 kD) is constitutively O-GlcNAcylated (**Figure 2A, 2B**). To rule out ambiguity due to N-glycosylation, we repeated the experiment in cell lysates pre-incubated with PNGase F removing this modification; O-GlcNAcylation was still observed (**Figure 2C**). Interestingly, no significant changes were observed when overall O-GlcNAcylation was increased with TMG. IP also identified a TRPC6-OGT protein interaction (**Figure 2D**). The O-GlcNAc status of TRPC6 was further explored using Click-it^TM^ in HEK-293T cells. O-GlcNAcylated protein residues were first labeled by tetra-acetylated azide-modified *N*-acetylglucosamine (GlcNAz) targeted to O-GlcNAc-modified residues by Y289L-b4-Gal-T1, and then co-tagged with biotin alkyne *via* click chemistry. The presence of biotinylation was determined by streptavidin-based immunoblot, and this assay also identified TRPC6 to be O-GlcNAcylated (**Figure 2E**).

**Figure 2.**
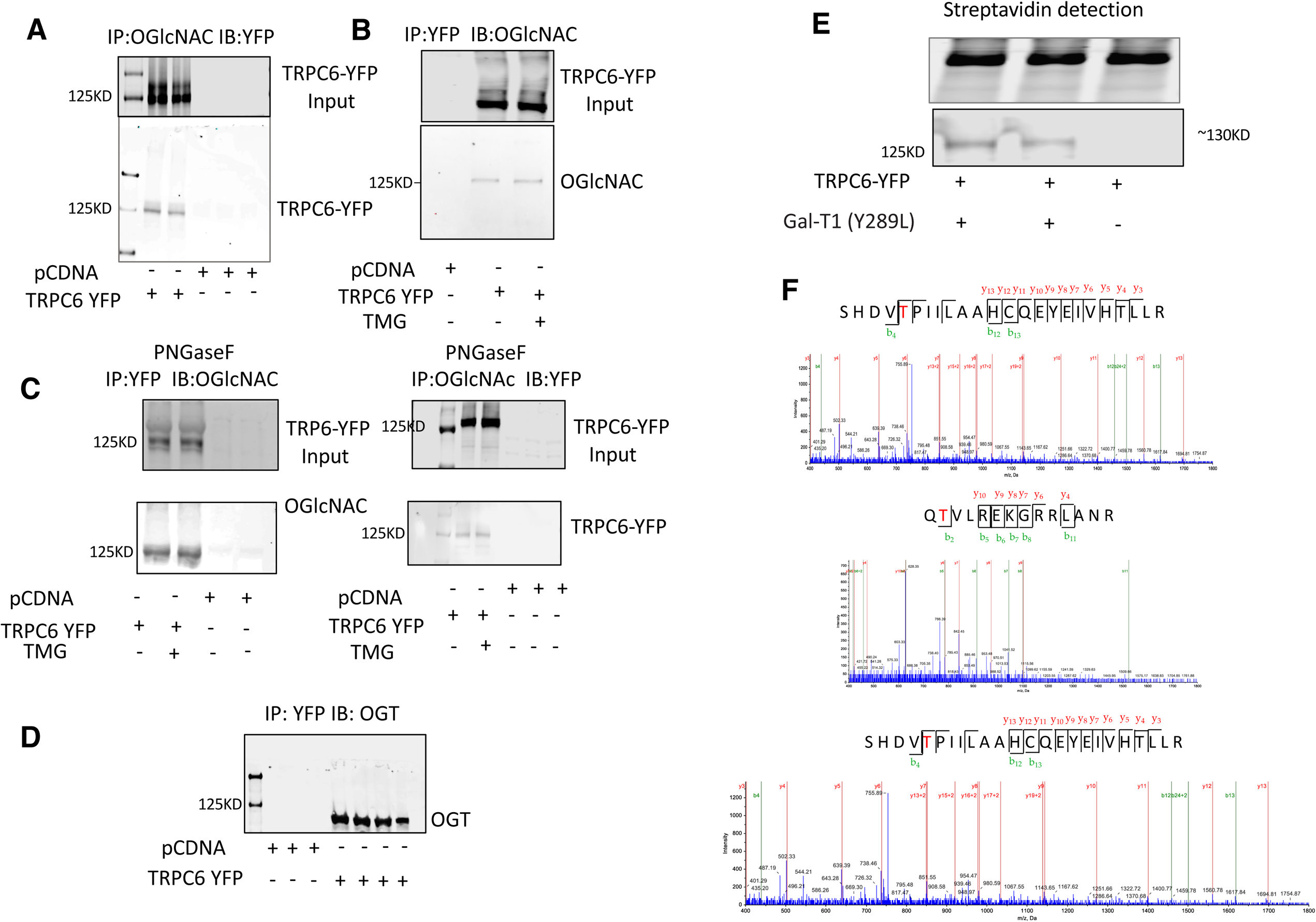
TRPC6 is constitutively OGlcNAcylated. **A)** HEK-293T expressing TRPC6-YFP (200 μg), lysate immunoprecipitated (IP) with O-GlcNAc antibody and immunoblotted (IB) with GFP antibody (detects YFP-labeled TRPC6). Upper gel input, lower IP. **B)** Same experiment with IP using YFP and IB for O-GlcNAc. Cells exposed to 10 μM Thiamet G for 24 h are also examined and show no change in IP bands. **C)** Representative IP blots with cells pre-treated with 0.2 µg of N-glycan-specific endoglycosidases PNGase F. **D)** HEK-293T cells co-transfected with TRPC6-YFP and OGT-Flag, IP using GFP Ab to capture TRPC6-YFP, and probed with OGT Ab. **E)** O-GlcNAc metabolic labeling of TRPC6-YFP by GlcNAz and mutated Gal-T1 to enable click chemistry that adds biotin to O-GlcNAcylated residues. Upper band shows input with IB for TRPC6-YFP, and lower band IP using GFP-Ab and IB with streptavidin to detect biotinylation. **F)** TRPC6-YFP generated in HEK-293T cells was immunoprecipitated with GFP Ab, and subjected to nano UPLC-MS/MS. MS/MS spectra identified O-GlcNAcylated peptides ‘RGS^14^SPRGAAGAAAR’,Q^70^TVLREKGRRLANR,‘SHDV^221^TPIILAAHC[CAM]QEYEIVHT LLR’ from the TRPC 6 protein. The position of O-GlcNAc sites are highlighted.

### T221 O-GlcNAcylation is a primary regulator of basal TRPC6 function

Based on these findings, we identified sites of TRPC6 O-GlcNAcylation by tandem mass spectrometry. Cell lysates from HEK-293T cells over-expressing TRPC6-YFP were subjected to pull-down assay using anti-YFP antibody magnetic beads. Immunoprecipitants were then subjected to in-gel digestion, and digests analyzed by nanoUPLC-MS/MS analysis. Three O-GlcNAc-containing peptides in TRPC6 were identified that mapped to S14, T70 and T221A (**Figure 2F**). Of these, T221 had not been previously reported to be modified by any PTM, whereas T70 was a known phosphorylation target of cGK1α and S14 of cdk5c(Liu *et al*., 2020b). T221 is in the AR4 domain of the TRPC6 channel and highly conserved across species (Supplemental Figure 1).

To investigate the functional roles of O-GlcNAcylated residues in TRPC6, we performed site-directed mutagenesis using alanine substitution at each site to prevent the PTM. Compared to wild-type (WT) TRPC6, S14A or T70A mutants modestly lowered basal NFAT activity by −19 and −36%, respectively (both P<0.002). By contrast, the T221A mutation resulted in a profound rise in NFAT activity (11-fold over WT, P<10^−12^, **Figure 3A**). Interestingly, when T221A was coexpressed with both S14A and T70A, NFAT activity doubled further (**Figure 3B**), revealing T221 as a dominant regulator of basal TRPC6-NFAT activity. To test if T221A disrupted membrane expression or external pore structure, we tested if a potent selective TRPC6 antagonist (BI 749327)(Lin *et al*., 2019) could still inhibit its NFAT activation. The inhibitor blocked both WT and T221A TRPC6 NFAT promotor activity markedly (**Figure 3C**). This indicates TRPC6-T221A membrane localization is intact whereas this declines if TRPC6 N-glycosylation is blocked(Talbot *et al*., 2019), and its exterior pore structure is likely unchanged.

**Figure 3.**
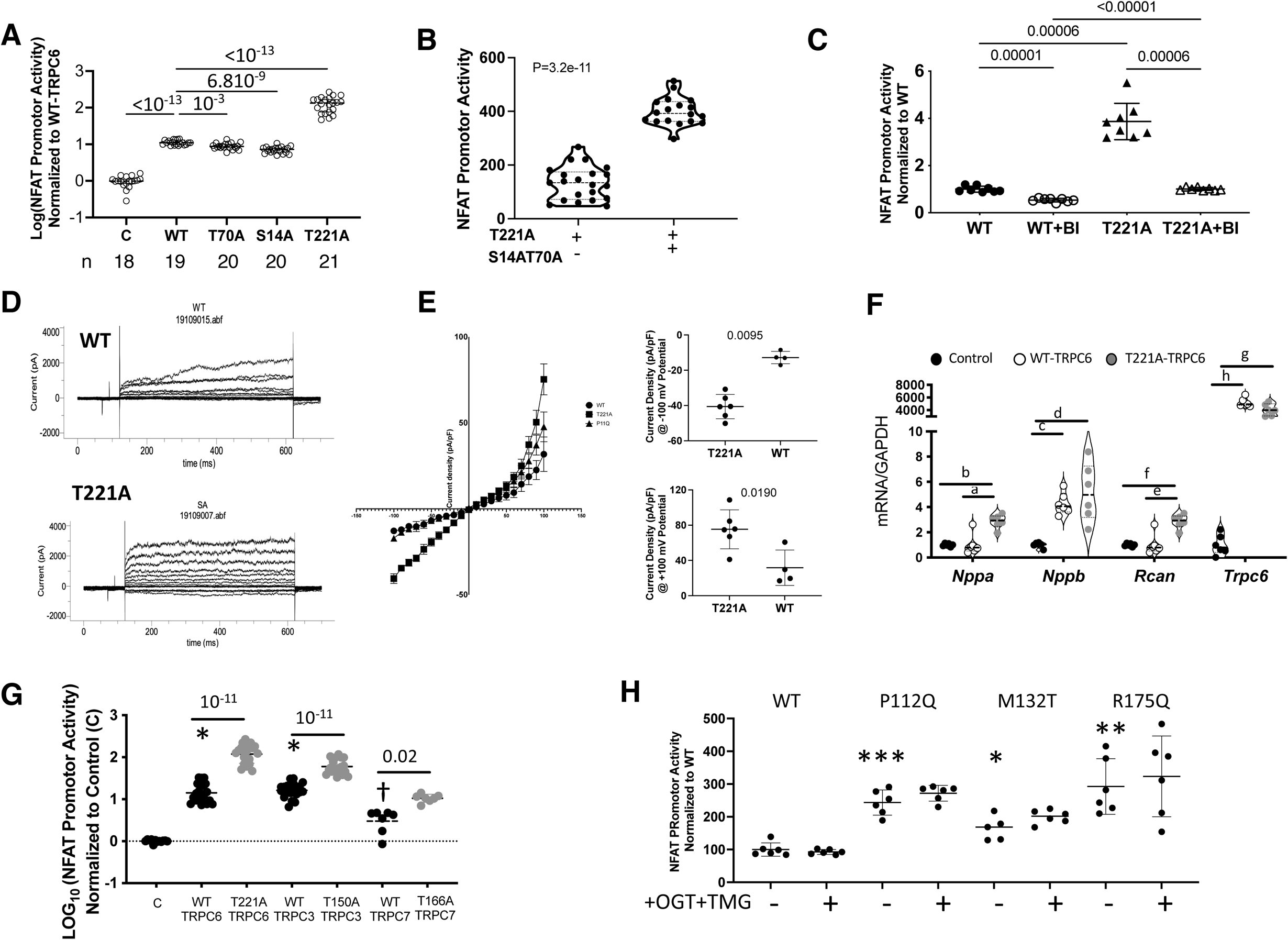
Substitution of T221 with T221A results in markedly hyMann WHitneyperactive TRPC6. **A)** NFAT promotor activity in cells transfected with pcDNA (C), TRPC6-YFP (WT) or TRPC6-YFP mutants (S14A, T70A, T221A) for 24 h (n/group provided in figure. Data log-transformed, Brown-Forsythe Welch ANOVA, Dunnett’s multiple comparison). **B)** Co-expression of *S*14A and T70A with T221A (n=18) versus T221A alone (n=21). P-value Mann-Whitney test. **C)** NFAT promotor activity due to expression of WT or T221A TRPC6 is similarly blunted by TRPC6 antagonist, BI 749327 (n=8/group, Welch ANOVA, Dunnett’s MCT.). **D)** Example current-time-voltage tracings in HEK-293 cells expressing WT or T221A mutant TRPC6. **E)** Summary current density vs voltage relations for WT, T221A, and P112Q TRPC6. Current density comparison at transmembrane voltage of +/− 100 mV is shown to right, P value Mann-Whitney. The current density plot also shows a gain-of-function mutation P112Q falls in between WT and T221A at positive voltages but has no impact on current density at negative voltages. **F)** Gene expression in rat neonatal cardiomyocytes expressing control vectors, TRPC6 wildtype (WT) or the T221A mutant for 48 hours. Genes are : A-type and B-type natriuretic peptide (*Nppa, Nppb),* regulator of calcineurin (*Rcan1*) and TRPC6 (*Trpc6*), each normalized to *Gapdh.* P-values displayed are from Dunnett’s MCT following Welch ANOVA; a – 0.0006, b – 0.00008, c – 0.00004, d – 0.005, e – 0.0006, f – 0.00008, g – 0.0003, h – 0.00002. (n=6 for *Trpc6*, n=7 for the rest). **G)** NFAT promotor activity (log-transformed) in control (C, n=13) and cells expressing either TRPC6 (WT vs T221A); TRPC3 (WT vs T150) (both n=21) and TRPC7 (WT vs T166) (n=7). P-values Dunnett’s MCT, Welch ANOVA. **H)** NFAT promotor activity in gain of function TRPC6 mutants +/− stimulated O-GlcNAcylation by combined TMG+OGT. The stimulation did not alter NFAT promotor activity for WT or any mutant. P-values for Mutant vs WT: * −0.033; ** 0.004; *** 0.0001 – Dunnett’s MCT/Welch ANOVA.

We next determined the functional impact of T221A. TRPC6 calcium conductance was measured using patch-clamp in HEK-293T cells. **Figure 3D** shows typical raw current-voltage-time tracings, and **Figure 3E** the summary data. Bi-directional voltage-dependent current increased by 75-80% in cells expressing T221A-TRPC6 versus WT. As a further comparator, we assessed this dependence in cells expressing P112Q-TRPC6, a potent gain of function (GOF) human mutation (Winn et al., 2005a). At positive voltages (outward current), the P122Q mutation was intermediate between WT and T221A, but at negative voltages (inward current), P122Q had no impact, whereas it was markedly increased by T221A (**Figure 3E**). These data identify T221 as a bi-directional regulator of basal TRPC6 Ca^2+^ current. Lastly, we expressed TRPC6 WT or T221A in cardiomyocytes finding T221A amplified gene expression coupled to pathological hypertrophy: *Nppa* (A-type natriuretic peptide)*, Nppb* (B-type natriuretic peptide)*, Myh7* (b-myosin heavy chain)*, Rcan1* (regulator of calcineurin-1), and *Trpc6* (**Figure 3F**).

Among the TRPC family of cation channels, TRPC3, TRPC6, and TRPC7 are the most homologous, the first two being very close in peptide sequence (Tang *et al*., 2018). Sequence analysis identified T150 in TRPC3 and T166 in TRPC7 as being homologous to T221 in TRPC6 (Supplemental Figure 2). We therefore mutated these sites to alanine to see if this modulation was also conserved. The results (**Figure 3G**) showed a similar marked elevation of basal NFAT promoter activity as a readout of channel conductance. This supports conservation of the regulatory role of this residue in the channel structure of all three TRPC channels.

Based on these data, we wondered if gain-of-function TRPC6 mutations that stimulate basal NFAT activity and reside in the N-terminus near T221 may work in part by interfering with T221 O-GlcNAcylation. This was tested in several potent GOF mutations each of which amplified resting NFAT promotor activity (**Figure 3H**) - though none to the level observed with the T221A mutation (e.g. 1-2x vs 11x). Furthermore, increasing O-GlcNAcylation with TMG (c.f. **Figure 1C**) had no impact (**Figure 3H**), indicating these GOF mutations do not likely interfere with T221 O-GlcNAcylation.

### Structural analysis reveals coordinated residues in AK4-linker region that impart basal TRPC6 regulation and O-GlcNAcylation control

To explore how T221-O-GlcNAcylation impacts TRPC6 channel function and identify if other coordinating amino acids are also involved, we turned to recent cryo-electron microscopy structural studies (Azumaya *et al*., 2018; Bai et al., 2020; Li et al., 2019) and modeled an O-GlcNAc modification to residue T221. The model structure of the relevant region is displayed in **Figure 4A** and predicts T221 O-GlcNAcylation (cyan) fosters electrostatic contacts between residues within the 193-203 loop in the AR4 domain crossing into the linker helix LH1 domain. The model also predicts O-GlcNAc at T221 enhances electrostatic interaction with glutamic acid (E246, purple) and hydrogen bond formation between the side chain of glutamine (Q198, pink) and the backbone amino group of aspartate (D205, light blue). We posited that these interactions hold a portion of the 193-203 loop in place to enhance S199 and E200 for interaction with O-GlcNAc at T221. The side chains of S199 (yellow), E200 (blue), and E246 are predicted to form hydrogen bonds with the hydroxyl of O-GlcNAc.

**Figure 4.**
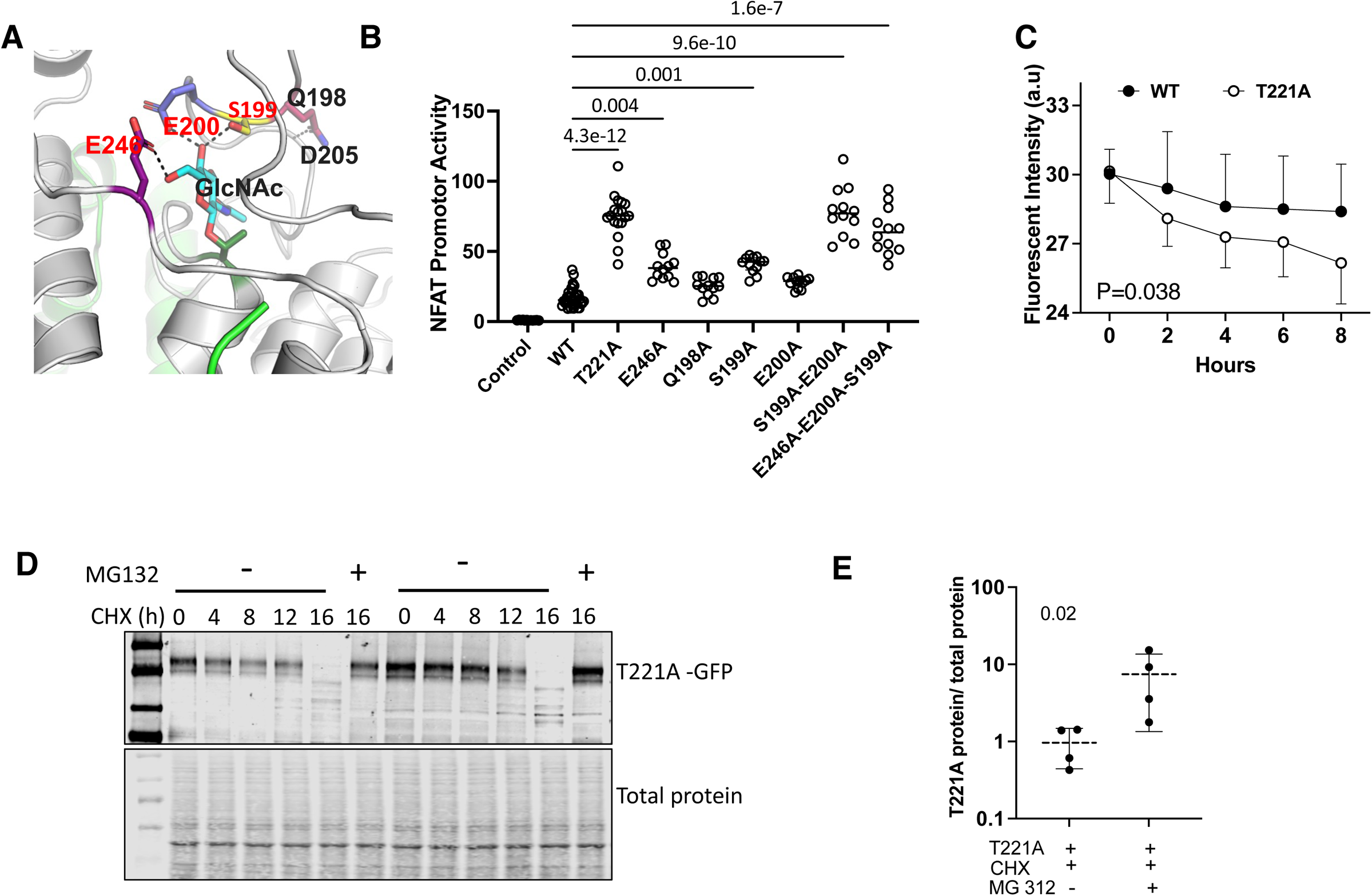
T221-coordinating amino acids in 192-203 Ankyrin Repeat-linker loop region are required for normal constrained TRPC6 channel function. **A)** Structural model of region linking O-GlcNAc modified T221 with AA193-203 loop near ankyrin repeat domain 4 and linker helix. Electrostatic interactions are between E246-*purple*, S199-*yellow*, E200-*blue*. **B)** HEK-293T cells transfected with pcDNA plasmid (Control), WT-TRPC6, and TRPC6 mutants impacting these predicted AA interactions. (n=33 for WT, 8 for T221A, and 12 for all other groups. Kruskal-Wallis test, Dunn’s MCT. **C)** Time dependent decline in WT (n=8) vs T221A (n=16) protein indexed by decline in YFP fluorescence in cells incubated with cycloheximide (CHX). Mean ± SD; P-value for slope difference by analysis of covariance. **D)** Western blot of experiment as depicted in panel C with protein expression of TRPC6-YFP shown in cells with or without co-treatment with MG132 to inhibit the proteasome. E) Densitometry of 16 hr post-CHX data +/− MG132. (n=4/group; Mann-Whitney test).

To test this, we selectively mutated each residue with alanine substitutions, expressed the recombinant forms of TRPC6 in HEK-293T cells, and assessed NFAT activity. The results (**Figure 4B**) show E246A and S199A mutations similarly increased NFAT activity over WT, but both less than with the T221A mutation. Neither Q198A nor E200A alone augmented activity over WT, however when both S199A and E200A were combined, the result was comparable with T221A. Adding E246A to form a triple mutant did not further increase NFAT activity. Thus, the coordinating hydrogen bonds between S199, E200, and O-GlcNAcylated T221 form critical residue interactions regulating basal TRPC6 conductance and NFAT-activation, with E246 itself also impacting this behavior.

### TRPC6 T221 O-GlcNAcylation enhances protein longevity

Beyond stimulating TRPC6 expression, channel conductance and NFAT signaling, we tested if T221 O-GlycNAcylation stabilizes the protein to protect it from proteasomal degradation. T221A or WT TRPC6 were expressed in HEK-293T cells exposed to cycloheximide to block de novo protein synthesis +/− the proteasome inhibitor MG132. The rate of protein decline was indexed by GFP fluorescence and was significantly faster with the T221A mutant (**Figure 4C**). This primarily reflected increased proteasome degradation as MG132 restored levels to baseline similarly with the mutant and WT form (**Figure 4D, 4E**). Thus, O-GlcNAcylation of TRPC6 reduces its proteasomal degradation to improve post-translational longevity.

## DISCUSSION

We identified O-GlcNAc residues in the N-terminal ankyrin repeat domain-4 of the TRPC6 channel, with T221 exerting primary control over basal channel current and associated NFAT activation. Preventing this modification results to our knowledge in the highest basal channel conductance and corresponding NFAT activity reported, including from multiple human GOF mutations causing disease or any other activation PTMs. Consistent with its augmentation of NFAT promoter activity, expression of T221A-TRPC6 amplifies a myocyte pathological hypertrophic gene program. We further find O-GlcNAcylation at T221 coordinates with several neighboring amino acids required to constrain channel conductance at normal levels and revealing this to be a key regulatory nexus. While we had first hypothesized that TRPC6 O-GlcNAcylation might be a mechanism for HG-stimulated TRPC6-dependent NFAT activation, our data refute this by showing that this modification is already maximal and inhibitory. The results identify a novel regulatory region of the protein that might be leveraged for therapeutics to constrain hyperactive TRPC6 and associated disease.

It is useful to frame our findings in the context of recent cryo-EM structural data regarding TRPC6 (Azumaya *et al*., 2018; Bai *et al*., 2020; Tang *et al*., 2018). Such studies reveal TRPC6 tetramers in a two-layered architecture assembled into an inverted bell-shaped intracellular cytosolic domain (ICD) that caps below the transmembrane domain (TMD). The ICD is assembled through interactions between the four ankyrin repeat domain (residues: 96–243) in the N-terminus, linker helices (residues: 256–393) and a coiled-coil domain in the C-terminus. ARs and LHs are key to inter-subunit interactions and TRPC6 tetramer assembly. Specifically, amino acids of the N-terminal loop (85–94) interact with LHs of the neighboring subunit and the last three LHs pack against the TRP helix providing the major contact site between the ICD and TMD. The ARs are highly conserved helix-turn-helix structural motifs (Bork, 1993; Li et al., 2006; Lux et al., 1990) involved in the assembly and stability of multiprotein complexes by forming both intra- and inter-repeat hydrophobic and hydrogen bond interactions (Erler et al., 2004; Mosavi et al., 2004). We find O-GlcNAcylated T221 forms stabilizing electrostatic contacts between AR4, the 193-203 loop near AR4, and loop connecting AR4 to LH1. We believe this helps hold distant regions together and is critical to stabilize the closed state of the channel pore. The six best known human GOF mutations that all cause renal disease are found in ARs: G109S (AR1), P112Q (AR1), N125S (AR1), M132T (AR2), N143S (AR2), R175Q(AR3) (Buscher et al., 2010; Gigante et al., 2011; Heeringa et al., 2009; Hofstra et al., 2013; Reiser *et al*., 2005; Santin *et al*., 2009; Winn *et al*., 2005b). Structural studies of these mutants similarly support destabilization of electrostatic interactions at the interface of AR domains and the linker helix that in turn associates with greater current (Bai *et al*., 2020). While our data did not suggest such mutations prevent T221 O-GlcNAcylation, it remains possible some others may, though unlikely fully or the impact on resting channel conductance would be too high. T221 mutations have not been reported in humans, and we suspect would be embryonic lethal. Still the location of the newly identified coordinated residue cluster T221-OGlcNAc, S199, E200, and E246 and Q198 confirms criticality of this conserved region and for constitutive O-GlcNAcylation.

While attention regarding O-GlcNAc modification has often been on its role in disease and potential as a modifiable PTM in that context, there is wide appreciation that it plays many constitutive roles (Chatham *et al*., 2021; Darley-Usmar et al., 2012; Wells et al., 2003). It is among the most abundant forms of protein glycosylation (Zachara et al., 2015), with OGT found in all metazoans and expressed in all mammalian tissues (Lubas et al., 1997; Watson et al., 2014). OGT also has non-catalytic functionality, but it appears only the O-GlcNAcylation function of OGT is required for cell survival (Levine et al., 2021). OGT strongly associates with the ribosome and nearly half of ribosomal proteins are O-GlcNAcylated (Zeidan et al., 2010). Furthermore, studies find O-GlcNAcylation of Sp1 and Nup62 proteins occur co-translationally and are key for their stability (Zhu et al., 2015). While a full list of OGT substrates obligatorily and constitutively modified for their cellular function and longevity remains to be assembled, and none have previously listed TRPC6, our data indicates this is a prime example. Interestingly, O-GlcNAcylation of phospholamban has been identified as a mechanism to impede S16 phosphorylation and in turn inhibit myocyte Ca^2+^ cycling into the sarcoplasmic reticulum (Yokoe et al., 2010). Another example is store-operated Ca^2+^ entry that is negatively controlled by STIM1 O-GlcNAcylation (Zhu-Mauldin et al., 2012), and could play roles in multiple mechanical and receptor-coupled signaling. O-GlcNAcylation of NFKB p65 and GSK-3β has been previously linked to reduced NFAT signaling with intermittent hypoxia (Nakagawa et al., 2019), but the sites of modification and constitutive status are not reported yet. While TRPC6 phophorylation bi-directionally modulates TRPC6 function and protein interaction, to our knowledge, this has not been found obligatory for basal functionality. This differs from N-glycosylation that is needed for channel membrane localization(Talbot *et al*., 2019) and as shown here O-GlcNAcylation needed to constrain channel conductance within the physiological range.

Our results remove O-GlcNAcylation of TRPC6 as a mechanism for *increased* activity with hyperglycemia but leave other mechanisms by which TRPC6 is stimulated such as oxidant stress and CAMKII activity augmented by O-GlcNAcylation, intact. They reveal a novel regulatory zone in TRPC6 essential for its normal function, and further highlight how O-GlcNAc modifications will require substantial nuancing if they are to have therapeutic value.

## Materials and Methods

### Plasmids

A pcDNA3-human TRPC6-YFP plasmid was obtained from Dr. Craig Montell (Koitabashi *et al*., 2010; Kwon et al., 2007), pcDNA3-human TRPC3 plasmid from Dr. Jeffery Molkentin (Seo *et al*., 2014b), pcDNA3-human TRPC7 from Dr. Steve S Pullen (Boehringer Ingelheim) and FLAG-OGT from Dr. Gerald W Hart. pGL4.30-NFAT-RE firefly luciferase (NFAT-luc) driven by the NFAT response element and Renilla luciferase (TK-Rluc) vectors were from Promega (Seo *et al*., 2014b). Alanine substitution mutants: TRPC6-YFP: S14A, T70A, T221A, S14AT221A, S14AT70AT221A, E246A, Q198A, S199A, E200A, S199AE200A, E246AS199AE200A, TRPC3-YFP: T150A; and TRPC7-YFP: T166A were each generated by PCR-based site mutagenesis (GeneArt Site-Directed Mutagenesis System, A13282) using pcDNA3-human TRPC6-YFP, TRPC3-YFP and TRPC7-YFP as the template. A human TRPC6 gene promoter construct was made by cloning a 1.7kb insert corresponding to the upstream of transcription start site of Human TRPC6, into luciferase reporter plasmid (pGL3 Luc, Promega U47295). A pcDNA3 vector served as the control for all transfection assays.

### HEK-293T cell transfection and luciferase promotor assay

HEK-293T cells were grown in Dulbecco’s modified Eagle’s medium supplemented with 10% fetal bovine serum, 2 mM glutamine, 1 mM pyruvate and antibiotics. Cells were cultured to 70% confluence and transfected with plasmids encoding NFAT-luc, TK-Rluc (internal control) and wildtype or alanine substituted TRPC mutants. Transfection was carried out using Xfect transfection reagent following manufacturer’s instruction (Takara Bio 631318). After transfection, cultures were maintained in serum containing medium for 24 h. All treatments were done in serum free culture medium. For High glucose (HG) exposure experiments, DMEM supplemented with 30 mM glucose (Sigma-Aldrich G8270) or 30 mM mannitol (Sigma-Aldrich M4125) (as HG control) was used. OGA inhibitor Thiamet G (TMG) (O-GlcNAc Core, JHU) was used at a concentration of 10 μM and glucosamine (GlcN) (Sigma-Aldrich G4875, 2 mM) was applied for 6h to stimulate O-GlcNAcylation (Chatham and Marchase, 2010). Cells were harvested using passive lysis buffer (Promega E1910). NFAT-luciferase activity was determined using Dual Luciferase Reporter Assay Kit (Promega E1910).

### Immunoprecipitation Assay

HEK-293T cells were transfected to express OGT-Flag and TRPC6-YFP. In some studies, cells were further treated with Thiamet G (TMG) to stimulate O-GlcNAcylation for 24h before harvesting. Cells were washed in ice-cold PBS (ThermoFisher Scientific 10010023), resuspended in RIPA buffer (Sigma-Aldrich R0278) containing Complete™ protease inhibitor cocktail (Roche Diagnostics, 11836153001) incubated for 20 min on ice to complete lysis. TMG 10 μM was also added to the lysis buffer in cells pre-treated with TMG. In some studies, PNGase F digestion (PNGase F, New England Biolabs P0704S) was done per manufacturer’s specifications to remove N-linked glycans. Cell lysates were centrifuged at 12000 rpm x 10min at 4°C and supernatants were used for immuno-precipitation (IP) analysis. IP was carried out by incubating lysates with OGT antibody (Sigma-Aldrich O6264), OGlcNAc antibody (CTD 110.6, MABS1254), or GFP M-trap beads (Chromotek GTD-20 serves as YFP capture beads) to capture TRPC6-YFP. 50 μl slurry of PierceTM Protein A/G Magnetic Beads (ThermoFisher Scientific 88802) were then added to OGT and OGlcNAc IP lysates and all IP samples were incubated for 4h at 4°C with gentle rolling. GFP M-trap beads and OGT IP magnetic beads were washed thrice with 1 ml of washing buffer (20 mM Tris/HCl, pH 7.4, 150 mM NaCl, 1 mM EDTA, 0.05% Triton X100, 5% glycerol and Complete™ protease inhibitor cocktail) and proteins were eluted in SDS-PAGE loading buffer. O-GlcNAc IP magnetic beads were washed 3x in TBS and eluted with 1M GlcNAc in TBS. Enriched proteins and 10% input samples were boiled in 30 μl of 2× SDS-PAGE loading buffer containing 5% 2-mercaptoethanol for 5 min, and further used for SDS-PAGE analysis.

### SDS/PAGE and Western blot analysis

For Western blot analysis, protein samples were separated by precast 4–15% Criterion™ TGX Stain-Free™ Protein Gels (Bio-Rad Laboratories 5678085), transferred to a nitrocellulose membrane using Trans-Blot® Turbo™ Midi Nitrocellulose Transfer Packs (Bio-Rad Laboratories 1704159EDU) and Trans-Blot® Turbo™ Transfer System (Bio-Rad Laboratories 1704150). The membrane was blocked with 5% non-fat dried skim milk solution for 1 h at 27°C, then incubated overnight at 4°C with primary antibodies diluted in blocking buffer: anti-OGT, anti-O-GlcNAc and anti-GFP (ThermoFisher Scientific A-6455). The next day, fluorescent secondary antibodies (IRDye 800CW donkey anti-rabbit 800, IRDye 800CW donkey anti-mouse 680, IRDye 800CW goat anti-rabbit 800 and IRDye 800CW goat anti-mouse 680 LI-COR), diluted in 1% milk in TBS-T (1L:L10L000), were added to the membranes for 1 h at room temperature. Images were acquired and analyzed using the LI-COR Odyssey Image System.

### O-GlcNAc metabolic labeling of WT-TRPC6

O-GlcNAcylation of TRPC6 was also assayed using Click-iT GlcNAz metabolic glycoprotein labeling reagents (ThermoFisher Scientific C33368). Lysates (200 μg protein) from HEK-293T cells expressing WT-TRPC6-YFP was incubated with GFP M-trap beads (Chromotek GTD-20) to immunoprecipitate TRPC6. Immunoprecipitated protein was enzymatically labeled utilizing the permissive mutant β-1,4-galactosyltransferase (Gal-T1 Y289L) which transfers azido-modified galactose (GalNAz) from UDP-GalNAz to O-GlcNAc residues on the target proteins (ThermoFisher Scientific C33368). The labelled lysate was then clicked on with biotin-alkyne using copper catalyzed azide-alkyne click chemistry reaction protocol according to manufacturer’s instruction (ThermoFisher Scientific C33372). The biotinylation was detected using IRDye 800CW Streptavidin dye and imaged (Odyssey Image System LICOR).

### nanoACQUITY UltraPerformance LC Mass Spectrometry

pcDNA3-human TRPC6-YFP was over-expressed in HEK-293T cells and immunoprecipitated. The eluate was subjected to SDS-PAGE followed by Commassie Blue staining. The corresponding gel bands were cut out and excised into cubes (ca. 1 × 1 mm) with a razor blade. Gel pieces were de-stained with 50% ACN followed by the addition of 100 ul of 10 mM dithiothreitol (DTT) in 50 mM bicarbonate buffer and incubation at 37°C for 0.5 h. After removal of DTT solution, 100 μl of 30 mM iodoacetamide in 50 mM bicarbonate buffer was added and incubated in dark for 30 minutes. Proteins were then digested with the addition of sequencing-grade trypsin/Lys-C followed by incubation at 37°C overnight. The yielded peptides were extracted and desalted with C18 Ziptip columns, with elutes dried down with a SpeedVac. Extracted peptides were analyzed with a NanoUPLC-MS/MS system integrating nanoAcquity UPLC (Waters) and a TripleTOF 6600 mass spectrometr (Sciex) (by using a similar setting as shown in a previous report (Aldeghaither et al., 2019), with some modifications. Specifically, dried peptides were dissolved in 0.1% formic acid and loaded onto a C18 Trap column (Waters Acquity UPLC Symmetry C18 NanoAcquity 10 K 2G V/M, 100 A, 5 μm, 180 μm x 20 mm) at 15 μl/min for 2 min. Peptides were then separated with an analytical column (Waters Acquity UPLC M-Class, peptide BEH C18 column, 300 A, 1.7 μm, 75 μm x 150 mm) which was temperature controlled at 40°C. The flow rate was set as 400 nL/min. A 60-min gradient of buffer A (2% ACN, 0.1% formic acid) and buffer B (0.1% formic acid in ACN) was used for separation: 1% buffer B at 0 min, 5% buffer B at 1 min, 45% buffer B at 35 min, 99% buffer B at 37min, 99% buffer B at 40 min, 1% buffer B at 40.1 min, and 1% buffer B at 60 min. Data were acquired with the TripleTOF 6600 mass spectrometer using an ion spray voltage of 2.3kV, GS1 5 psi, GS2 0, CUR 30 psi and an interface heater temperature of 150°C. Mass spectra was recorded with Analyst TF 1.7 software in the IDA mode. Each cycle consisted of a full scan (m/z 400-1600) and fifty information dependent acquisitions (IDAs) (m/z 100-1800) in the high sensitivity mode with a 2+ to 5+ charge state. Rolling collision energy was used.

Data files were submitted for simultaneous searches using Protein Pilot version 5.0 software (Sciex) utilizing the Paragon and Progroup algortihms and the integrated false discovery rate (FDR) analysis function. MS/MS data was searched against the customized human TRPC6 protein database. Trypsin/LysC was selected as the enzyme. Carbamidomethylation was set as a fixed modification on cysteine. HexNAc emphasis was chosen as a special factor. Other search parameters include instrument (TripleTOF 6600), ID Focus (Biological modifications), search effort (Thorough), false discovery rate (FDR) analysis (Yes), and user modified parameter files (No). The proteins were inferred based on the ProGroupTM algorithm using ProteinPilot software. The detected protein threshold in the software was set to the value which corresponded to 5% FDR. Peptides were defined as redundant if they had identical cleavage site(s), amino acid sequence, and modification. All peptides were filtered with confidence to 5% FDR, with the confidence of HexNAc sites automatically calculated. Each of the HexNAc modification sites (>95% confidence) was then manually confirmed and annotated (Ma and Hart, 2017).

### Electrophysiology studies – Patch Clamp

Patch clamp studies were done as previously described protocol (Hall et al., 2014). Briefly, HEK-293T cells were transfected with 1 µg of TRPC6 channel plasmids (WT, T221A and 112Q) using Xfect transfection reagent according to manufacturer’s protocol (Takara Bio 631318). Cells expressing wild type and mutant channels were identified by YFP fluorescence. The bath solution was 140mM NaCl, 5mM CsCl2, 1 mM MgCl2, 10 mM HEPES, and 10 mM glucose with pH of 7.4. Borosilicate glass capillary pipettes (World Precision Instr.) were used with ~3MΩ resistance when filled with solution containing 5mM NaCl, 40 mM CsCl2, 80mM Cs-glutamate, 5mM Mg-ATP, 5 mM EGTA, 1.5 CaCl2 (free calcium concentration was 100 nM). Currents were obtained using a voltage step-pulse protocol or ramp protocol from −100 mV to +100 mV applied every 2s for 500 ms from holding potential of −60 mV. Current recordings were in a whole-cell configuration using Axopatch 200A amplifier (Axon Instruments, Molecular Devices). Data are represented as mean standard error of mean, and analyzed by One-Way ANOVA or by Two-way ANOVA with Tukey multiple comparisons test.

### Neonatal Rat Ventricular myocyte isolation and adenoviral transfection

Neonatal Rat cardiomyocytes (NRVMs) were freshly isolated as previously described (Mishra et al., 2021) and cultured at 1 million cells per well in six-well plates for 24 hr in DMEM with 10% FBS and 1% penicillin/streptomycin. Adenoviruses were developed expressing the GFP-tagged wild-type sequence of human TRPC6 or TRPC6 T221A. NRVMs were infected with an MOI of 10 with the respective viruses for 48 hr before performing the downstream assays.

### RNA isolation and Gene expression analysis

Total RNA from NRVMs was extracted using Trizol Reagent (Cat. No. 15596026, Invitrogen, Thermofisher, USA) per manufacturer’s instructions. High-Capacity RNA-to-cDNA Kit (Cat. No. 4388950, Applied Biosystems, Thermofisher, USA) was used to reverse transcribe the RNA into cDNA as described before (Mishra *et al*., 2021). Quantitative real time PCR analysis was carried out using TaqMan specific primers for: *Trpc6* (Rn00677559_m1), *Nppa* (Rn00664637_g1), *Nppb* (Rn00580641_m1), *Myh7* (Rn01488777_g1), and *Rcan1* (Rn01458494_m1), and *Gapdh* (Rn01775763_g1) from Applied Biosystems. The threshold cycle (Ct) values were determined by crossing point method and normalized to GAPDH (Applied Biosystems) values for each run.

### Protein degradation measurement

A fluorescent microplate-based assay was used to measure TRPC6-WT versus TRPC6-T221A-GFP signal intensity decay. HEK-293T cells seeded on 96 well plates were transfected with 0.1 μg of the respective plasmids. 24 h after transfection, cells were treated with cycloheximide (100 μg/ml), and fluorescence intensities of the wells were measured at the indicated time points. T221A-GFP decline was measured in cycloheximide treated cells in the presence or absence of proteasome inhibitor MG132 (10 μM). For western blot experiments, 24 h after transfection, HEK cells were incubated with 100 μg/ml of cycloheximide. Cells were harvested at the indicated time points as shown in Fig 4c, followed by cell lysis, SDS-PAGE and western blotting to visualize TRPC6 T221A protein levels.

### Statistical Analysis

Statistical analysis was performed using Prism Ver 9.3.1. All of the individual tests used for each data figure in the study are provided in their respective figure legend along with sample size per group. For multiple groups, 1-way ANOVA, a Welch ANOVA (if test for variance difference between groups was positive), or non-parametric Kruskal Wallis (if non-normally distributed) was used. A 2-Way ANOVA was also used in some testing as indicated. Two-group comparisons used non-parametric tests (Mann Whitney U test). All precise p values are provided for statistical testing in the figures and/or legends.

## Acknowledgements

This study was supported by National Institute of Health – Heart Lung and Blood Institute grants (R35 HL135827 (DAK), P01HL107153 (DAK, SM), and General Medicine (R35GM128595, RCP), American Heart Association Grants (CDA938718 (SM), and CDA34110140 (MJR).

## Author Contributions

SM performed the majority of the experiments, data analysis, figure and manuscript preparation; JM performed the mass spectroscopy studies; MS generated the site mutation plasmid vectors, FF performed the patch clamp electrophysiologic studies and analysis; RCP performed the molecular structural analysis and interacting residue predictions; MJR assisted with molecular assays and editorial input into the manuscript; NZ provided expertise in O-GlcNAc biology and experimental design and interpretation; and DAK supervised the project, provided input into study design, data analysis, and edited the manuscript.

## Declaration of Interests

DAK receives grant support from Boehringer Ingelheim pursuing the use of a selective TRPC6 antagonist for treatment of Duchenne Muscular Dystrophy. None of the other authors declare any competing interests.

## Notes

### Competing Interest Statement

The authors have declared no competing interest.

