## Supplemental Figures for "Constitutive Inhibition of Transient Receptor Potential Canonical Type 6 (TRPC6) by O-GlcNAcylation at Threonine-221"

Supplemental Figure 1: Amino acid sequence in AK4-LK1 region of TRPC6 where T221 and coordinating amino acids reside is highly conserved across many species.

NCBI Multiple Sequence Alignment Viewer, Version 1.22.0

| Sequence ID | | Start | Alignment | End | Organism |
|---|---|---|---|---|---|
| | | | 200 210 220 230 240 250 | | |
| NP_004612.2 | (+) | 1 | E L Q Q D D F Y A Y D E D G T R F S H D V T P I I L A A H C Q E Y E I V H T L L R K G A R I E R P H D | 931 | Homo sapiens |
| NP_038866.2 | (+) | 1 | E L Q Q D D F Y A Y D E D G T R F S H D V T P I I L A A H C Q E Y E I V H T L L R K G A R I E R P H D | 930 | Mus musculus |
| NP_446011.1 | (+) | 1 | E L Q Q D D F Y A Y D E D G T R F S H D V T P I I L A A H C Q E Y E I V H T L L R K G A R I E R P H D | 930 | Rattus norvegicus |
| XP_003357304.3 | (+) | 1 | E L Q Q D D F Y A Y D E D G T R F S H D V T P I I L A A H C Q E F E I V H T L L R K G A R I E R P H D | 931 | Sus scrofa |
| XP_014971021.2 | (+) | 1 | E L Q Q D D F Y A Y D E D G T R F S H D V T P I I L A A H C Q E Y E I V H T L L R K G A R I E R P H D | 931 | Macaca mulatta |
| XP_546553.4 | (+) | 1 | E F Q Q D D F Y A Y D E D G T R F S H D V T P I I L A A H C Q E Y E I V H T L L R K G A R I E R P H D | 932 | Canis lupus familiaris |
| XP_016777341.2 | (+) | 1 | E L Q Q D D F Y A Y D E D G T R F S H D V T P I I L A A H C Q E Y E I V H T L L R K G A R I E R P H D | 931 | Pan troglodytes |
| NP_001166503.1 | (+) | 1 | E L Q Q D D F Y A Y D E D G T R F S H D V T P I I L A A H C Q E Y E I V H T L L R K G A R I E R P H D | 893 | Cavia porcellus |
| XP_034521520.1 | (+) | 1 | E L Q Q D D F Y A Y D E D G T R F S H D V T P I I L A A H C Q E Y E I V H T L L R K G A R I E R P H D | 960 | Ailuropoda melanoleuca |
| XP_032344153.1 | (+) | 1 | E L Q Q D D F Y A Y D E D G T R F S H D V T P I I L A A H C Q E Y E I V H T L L R K G A R I E R P H D | 932 | Camelus ferus |
| XP_012917898.1 | (+) | 1 | E L Q Q D D F Y A Y D E D G T R F S H D V T P I I L A A H C Q E Y E I V H T L L R K G A R I E R P H D | 883 | Mustela putorius furo |
| XP_014334446.1 | (+) | 1 | E L Q Q D D F Y A Y D E D G T R F S H D V T P I I L A A H C Q E Y E I V H T L L R K G A R I E R P H D | 846 | Bos mutus |
| XP_005077380.2 | (+) | 1 | E L Q Q D D F Y A Y D E D G T R F S H D V T P I I L A A H C Q E Y E I V H T L L R K G A R I E R P H D | 929 | Mesocricetus auratus |
| XP_001499671.2 | (+) | 1 | E L Q Q D D F Y A Y D E D G T R F S H D V T P I I L A A H C Q E Y E I V H T L L R K G A R I E R P H D | 932 | Equus caballus |
| XP_002754689.1 | (+) | 1 | E L Q Q D D F Y A Y D E D G T R F S H D V T P I I L A A H C Q E Y E I V H T L L R K G A R I E R P H D | 931 | Callithrix jacchus |
| XP_040833147.1 | (+) | 1 | E L Q Q D D F Y A Y D E D G T R F S H D V T P I I L A A H C Q E Y E I V H T L L R K G A R I E R P H D | 929 | Ochotona curzoniae |
| XP_004585062.1 | (+) | 1 | E L Q Q D D F Y A Y D E D G T R F S H D V T P I I L A A H C Q E Y E I V H T L L R K G A R I E R P H D | 929 | Ochotona princeps |
| XP_002708649.1 | (+) | 1 | E L Q L D D F Y A Y D E D G T R F S H D V T P I I L A A H C Q E Y E I V H T L L R K G A R I E R P H D | 929 | Oryctolagus cuniculus |
| XP_020017264.1 | (+) | 1 | E L Q Q D D F Y A Y D E D G T R F S H D V T P I I L A A H C Q E Y E I V H T L L R K G A R I E R P H D | 931 | Castor canadensis |
| XP_021027930.1 | (+) | 1 | E L Q Q D D F Y A Y D E D G T R F S H D V T P I I L A A H C Q E Y E I V H T L L R K G A R I E R P H D | 930 | Mus caroli |
| XP_021062063.1 | (+) | 1 | E L Q Q D D F Y A Y D E D G T R F S H D V T P I I L A A H C Q E Y E I V H T L L R K G A R I E R P H D | 930 | Mus pahari |
| XP_006983838.1 | (+) | 1 | E L Q Q D D F Y A Y D E D G T R F S H D V T P I I L A A H C Q E Y E I V H T L L R K G A R I E R P H D | 927 | Peromyscus maniculat... |
| XP_004661969.1 | (+) | 1 | E L Q Q D D F Y A Y D E D G T R F S H D V T P I I L A A H C Q E Y E I V H T L L R K G A R I E R P H D | 931 | Jaculus jaculus |
| XP_021483360.1 | (+) | 1 | E L Q Q D D F Y A Y D E D G T R F S H D V T P I I L A A H C Q E Y E I V H T L L R K G A R I E R P H D | 845 | Meriones unguiculatus |
| XP_032766559.1 | (+) | 1 | E L Q Q D D F Y A Y D E D G T R F S H D V T P I I L A A H C Q E Y E I V H T L L R K G A R I E R P H D | 930 | Rattus rattus |
| XP_028619194.1 | (+) | 1 | E L Q Q D D F Y A Y D E D G T R F S H D V T P I I L A A H C Q E Y E I V H T L L R K G A R I E R P H D | 930 | Grammomys surdaster |
| XP_031200579.1 | (+) | 1 | E L Q Q D D F Y A Y D E D G T R F S H D V T P I I L A A H C Q E Y E I V H T L L R K G A R I E R P H D | 930 | Mastomys coucha |
| XP_034347500.1 | (+) | 1 | E L Q Q D D F Y A Y D E D G T R F S H D V T P I I L A A H C Q E Y E I V H T L L R K G A R I E R P H D | 930 | Arvicanthis niloticus |
| XP_003496186.1 | (+) | 1 | E L Q Q D D F Y A Y D E D G T R F S H D V T P I I L A A H C Q E Y E I V H T L L R K G A R I E R P H D | 929 | Cricetulus griseus |
| XP_028746785.1 | (+) | 1 | E L Q Q D D F Y A Y D E D G T R F S H D V T P I I L A A H C Q E Y E I V H T L L R K G A R I E R P H D | 929 | Peromyscus leucopus |
| XP_036050251.1 | (+) | 1 | E L Q Q D D F Y A Y D E D G T R F S H D V T P I I L A A H C Q E Y E I V H T L L R K G A R I E R P H D | 929 | Onychomys torridus |
| XP_038178700.1 | (+) | 1 | E L Q Q D D F Y A Y D E D G T R F S H D V T P I I L A A H C Q E Y E I V H T L L R K G A R I E R P H D | 929 | Arvicola amphibius |
| XP_041513139.1 | (+) | 1 | E L Q Q D D F Y A Y D E D G T R F S H D V T P I I L A A H C Q E Y E I V H T L L R K G A R I E R P H D | 929 | Microtus oregoni |
| XP_005346846.1 | (+) | 1 | E L Q Q D D F Y A Y D E D G T R F S H D V T P I I L A A H C Q E Y E I V H T L L R K G A R I E R P H D | 929 | Microtus ochrogaster |
| XP_013366708.1 | (+) | 1 | E L Q Q D D F Y A Y D E D G T R F S H D V T P V I L A A H C Q E Y E I V H T L L R K G A R I E R P H D | 929 | Chinchilla lanigera |
| XP_004626213.1 | (+) | 1 | E L Q H D D F Y A Y D E D G T R F S H D V T P I I L A A H C Q E Y E I V H T L L R K G A R I E R P H D | 845 | Octodon degus |
| XP_004870862.1 | (+) | 1 | E L Q Q D D F Y A Y D E D G T R F S H D V T P I I L A A H C Q E Y E I V H T L L R K G A R I E R P H D | 931 | Heterocephalus glaber |
| XP_010640339.1 | (+) | 1 | E L Q Q D D F Y A Y D E D G T R F S H D V T P I I L A A H C Q E Y E I V H T L L R K G A R I E R P H D | 931 | Fukomys damarensis |
| XP_013215690.1 | (+) | 1 | E L Q Q D D F Y A Y D E D G T R F S H D V T P I I L A A H C Q E Y E I V H T L L R K G A R I E R P H D | 911 | Ictidomys tridecemlineatus |
| XP_027786504.1 | (+) | 1 | E L Q Q D D F Y A Y D E D G T R F S H D V T P I I L A A H C Q E Y E I V H T L L R K G A R I E R P H D | 931 | Marmota flaviventris |
| XP_041625852.1 | (+) | 1 | E L Q Q D D F Y A Y D E D G T R F S H D V T P I I L A A H C Q E Y E I V H T L L R K G A R I E R P H D | 932 | Vulpes lagopus |
| XP_025862526.1 | (+) | 1 | E L Q Q D D F Y A Y D E D G T R F S H D V T P I I L A A H C Q E Y E I V H T L L R K G A R I E R P H D | 846 | Vulpes vulpes |
| XP_040486594.1 | (+) | 1 | E L Q Q D D F Y A Y D E D G T R F S H D V T P I I L A A H C Q E Y E I V H T L L R K G A R I E R P H D | 906 | Ursus maritimus |
| XP_025749691.1 | (+) | 1 | E L Q Q D D F Y A Y D E D G T R F S H D V T P I I L A A H C Q E Y E I V H T L L R K G A R I E R P H D | 932 | Callorhinus ursinus |
| XP_027961810.1 | (+) | 1 | E L Q Q D D F Y A Y D E D G T R F S H D V T P I I L A A H C Q E Y E I V H T L L R K G A R I E R P H D | 932 | Eumetopias jubatus |
| XP_027435468.1 | (+) | 1 | E L Q Q D D F Y A Y D E D G T R F S H D V T P I I L A A H C Q E Y E I V H T L L R K G A R I E R P H D | 932 | Zalophus californianus |
| XP_021552267.1 | (+) | 1 | E L Q Q D D F Y A Y D E D G T R F S H D V T P I I L A A H C Q E Y E I V H T L L R K G A R I E R P H D | 504 | Neomonachus schauins... |
| XP_035971783.1 | (+) | 1 | E L Q Q D D F Y A Y D E D G T R F S H D V T P I I L A A H C Q E Y E I V H T L L R K G A R I E R P H D | 932 | Halichoerus grypus |
| XP_006749681.1 | (+) | 1 | E L Q Q D D F Y A Y D E D G T R F S H D V T P I I L A A H C Q E Y E I V H T L L R K G A R I E R P H D | 932 | Leptonychotes weddellii |
| XP_034864658.1 | (+) | 1 | E L Q Q D D F Y A Y D E D G T R F S H D V T P I I L A A H C Q E Y E I V H T L L R K G A R I E R P H D | 932 | Mirounga leonina |
| XP_032283806.1 | (+) | 1 | E L Q Q D D F Y A Y D E D G T R F S H D V T P I I L A A H C Q E Y E I V H T L L R K G A R I E R P H D | 932 | Phoca vitulina |
| XP_010846532.1 | (+) | 1 | E L Q Q D D F Y A Y D E D G T R F S H D V T P I I L A A H C Q E Y E I V H T L L R K G A R I E R P H D | 877 | Bison bison bison |
| XP_019831324.1 | (+) | 1 | E L Q Q D D F Y A Y D E D G T R F S H D V T P I I L A A H C Q E Y E I V H T L L R K G A R I E R P H D | 846 | Bos indicus |
| XP_024616129.1 | (+) | 1 | E L Q Q D D F Y A Y D E D G T R F S H D V T P I V L A A H C Q E Y E I V H T L L R K G A R I E R P H D | 932 | Neophocaena asiaeorie... |
| XP_007191273.1 | (+) | 1 | E L Q Q D D F Y A Y D E D G T R F S H D V T P I V L A A H C Q E Y E I V H T L L R K G A R I E R P H D | 932 | Balaenoptera acutorostr... |
| XP_007448608.1 | (+) | 1 | E L Q Q D D F Y A Y D E D G T R F S H D V T P I V L A A H C Q E Y E I V H T L L R K G A R I E R P H D | 932 | Lipotes vexillifer |
| XP_026943551.1 | (+) | 1 | E L Q Q D D F Y A Y D E D G T R F S H D V T P I V L A A H C Q E Y E I V H T L L R K G A R I E R P H D | 932 | Lagenorhynchus obliqui... |
| XP_030739561.1 | (+) | 1 | E L Q Q D D F Y A Y D E D G T R F S H D V T P I V L A A H C Q E Y E I V H T L L R K G A R I E R P H D | 932 | Globicephala melas |
| XP_004283396.1 | (+) | 1 | E L Q Q D D F Y A Y D E D G T R F S H D V T P I V L A A H C Q E Y E I V H T L L R K G A R I E R P H D | 932 | Orcinus orca |
| XP_033718002.1 | (+) | 1 | E L Q Q D D F Y A Y D E D G T R F S H D V T P I V L A A H C Q E Y E I V H T L L R K G A R I E R P H D | 932 | Tursiops truncatus |
| XP_032496065.1 | (+) | 1 | E L Q Q D D F Y A Y D E D G T R F S H D V T P I V L A A H C Q E Y E I V H T L L R K G A R I E R P H D | 932 | Phocoena sinus |
| XP_029064752.1 | (+) | 1 | E L Q Q D D F Y A Y D E D G T R F S H D V T P I V L A A H C Q E Y E I V H T L L R K G A R I E R P H D | 932 | Monodon monoceros |
| XP_022417762.1 | (+) | 1 | E L Q Q D D F Y A Y D E D G T R F S H D V T P I V L A A H C Q E Y E I V H T L L R K G A R I E R P H D | 932 | Delphinapterus leucas |
| XP_023971087.1 | (+) | 1 | E L Q Q D D F Y A Y D E D G T R F S H D V T P I V L A A H C Q E Y E I V H T L L R K G A R I E R P H D | 932 | Physeter catodon |
| XP_036718287.1 | (+) | 1 | E L Q Q D D F Y A Y D E D G T R F S H D V T P I V L A A H C Q E Y E I V H T L L R K G A R I E R P H D | 932 | Balaenoptera musculus |
| XP_007520782.1 | (+) | 1 | E L Q H D D F Y A Y D E D G T R F S H D V T P I I L A A H C Q E Y E I V H T L L R K G A R I E R P H D | 928 | Erinaceus europaeus |
| XP_004604872.1 | (+) | 1 | E L Q Q D D F Y A Y D E D G T R F S H D V T P I I L A S H C Q E Y E I V H T L L R K G A R I E R P H D | 846 | Sorex araneus |
| XP_004689485.2 | (+) | 1 | E L Q L D D F Y A Y D E D G T R F S H D V T P V I L A A H C Q E Y E I V H T L L R R G A R I E R P H D | 876 | Condylura cristata |
| XP_037360681.1 | (+) | 1 | E L Q L D D F Y A Y D E D G T R F S H D V T P I I L A A H C Q E Y E I V H T L L R K G A R I E R P H D | 846 | Talpa occidentalis |
| XP_011367719.1 | (+) | 1 | E L Q Q D D F Y A Y D E D G T R F S H D V T P I V L A A H C Q E Y E I V H T L L R K G A R I E R P H D | 932 | Pteropus vampyrus |
| XP_039737901.1 | (+) | 1 | E L Q Q D D F Y A Y D E D G T R F S H D V T P I V L A A H C Q E Y E I V H T L L R K G A R I E R P H D | 932 | Pteropus giganteus |
| XP_006907565.1 | (+) | 1 | E L Q Q D D F Y A Y D E D G T R F S H D V T P I V L A A H C Q E Y E I V H T L L R K G A R I E R P H D | 932 | Pteropus alecto |
| XP_015989491.2 | (+) | 1 | E L Q Q D D F Y A Y D E D G T R F S H D V T P I V L A A H C Q E Y E I V H T L L R K G A R I E R P H D | 932 | Rousettus aegyptiacus |
| XP_032974848.1 | (+) | 1 | E L Q Q D D F Y A Y D E D G T R F S H D V T P I I L A A Q C Q E Y E I V H T L L R K G A R I E R P H D | 932 | Rhinolophus ferrumequin... |
| XP_037012893.1 | (+) | 1 | E L Q Q D D F Y A Y D E D G T R F S H D V T P V I L A A H C Q E Y E I V H T L L R K G A R I E R P H D | 932 | Artibeus jamaicensis |
| XP_024430371.1 | (+) | 1 | E L Q Q D D F Y A Y D E D G T R F S H D V T P V I L A A H C Q E Y E I V H T L L R K G A R I E R P H D | 932 | Desmodus rotundus |
| XP_028373336.1 | (+) | 1 | E L Q Q D D F Y A Y D E D G T R F S H D V T P I I L A A H C Q E Y E I V H T L L R K G A R I E R P H D | 932 | Phyllostomus discolor |
| XP_036282984.1 | (+) | 1 | E L Q Q D D F Y A Y D E D G T R F S H D V T P I I L A A H C Q E Y E I V H T L L R K G A R I E R P H D | 931 | Pipistrellus kuhlii |
| XP_008147418.1 | (+) | 1 | E L Q Q D D F Y A Y D E D G T R F S H D V T P V I L A A H C Q E Y E I V H T L L R K G A R I E R P H D | 932 | Eptesicus fuscus |
| XP_005872271.1 | (+) | 1 | E L Q Q D D F Y A Y D E D G T R F S H D V T P V I L A A H C Q E Y E I V H T L L R K G A R I E R P H D | 871 | Myotis brandtii |
| XP_006775322.2 | (+) | 1 | E L Q Q D D F Y A Y D E D G T R F S H D V T P V I L A A H C Q E Y E I V H T L L R K G A R I E R P H D | 814 | Myotis davidii |
| XP_036180497.1 | (+) | 1 | E L Q Q D D F Y A Y D E D G T R F S H D V T P V I L A A H C Q E Y E I V H T L L R K G A R I E R P H D | 926 | Myotis myotis |
| XP_023614611.1 | (+) | 1 | E L Q Q D D F Y A Y D E D G T R F S H D V T P V I L A A H C Q E Y E I V H T L L R K G A R I E R P H D | 856 | Myotis lucifugus |
| XP_036114295.1 | (+) | 1 | E L Q Q D D F Y A Y D E D G T R F S H D V T P V I L A A H C Q E Y E I V H T L L R K G A R I E R P H D | 932 | Molossus molossus |
| XP_014718249.1 | (+) | 1 | E L Q Q D D F Y A Y D E D G T R F S H D V T P I I L A A H C Q E Y E I V H T L L R K G A R I E R P H D | 932 | Equus asinus |
| XP_008529351.1 | (+) | 1 | E L Q Q D D F Y A Y D E D G T R F S H D V T P I I L A A H C Q E Y E I V H T L L R K G A R I E R P H D | 932 | Equus przewalskii |
| XP_036758704.1 | (+) | 1 | E L Q L D D F Y A Y D E D G T R F S H D V T P I I L A A H C Q E Y E I V H T L L R K G A R I E R P H D | 932 | Manis pentadactyla |
| XP_036881527.1 | (+) | 1 | E L Q L D E F Y A Y D E D G T R F S H D V T P I I L A A H C Q E Y E I V H T L L R K G A R I E R P H D | 932 | Manis javanica |
| XP_008588083.1 | (+) | 1 | E L Q Q D D F Y A Y D E D G T R F S H D V T P I I L A A H C Q E Y E I V H T L L R K G A R I E R P H D | 876 | Galeopterus variegatus |
| XP_027628284.1 | (+) | 1 | E L Q Q D D F Y A Y D E D G T R F S H D V T P I I L A A H C Q E Y E I V H T L L R K G A R I E R P H D | 930 | Tupaia chinensis |
| XP_011782001.1 | (+) | 1 | E L Q Q D D F Y A Y D E D G T R F S H D V T P I I L A A H C Q E Y E I V H T L L R K G A R I E R P H D | 931 | Colobus angolensis pall... |
| XP_037861213.1 | (+) | 1 | E L Q Q D D F Y A Y D E D G T R F S H D V T P I I L A A H C Q E Y E I V H T L L R K G A R I E R P H D | 931 | Chlorocebus sabaeus |
| XP_011927452.1 | (+) | 1 | E L Q Q D D F Y A Y D E D G T R F S H D V T P I I L A A H C Q E Y E I V H T L L R K G A R I E R P H D | 959 | Cercocebus atys |
| XP_005579492.1 | (+) | 1 | E L Q Q D D F Y A Y D E D G T R F S H D V T P I I L A A H C Q E Y E I V H T L L R K G A R I E R P H D | 931 | Macaca fascicularis |
| XP_011713765.1 | (+) | 1 | E L Q Q D D F Y A Y D E D G T R F S H D V T P I I L A A H C Q E Y E I V H T L L R K G A R I E R P H D | 931 | Macaca nemestrina |
| XP_031508878.1 | (+) | 1 | E L Q Q D D F Y A Y D E D G T R F S H D V T P V I L A A H C Q E Y E I V H T L L R K G A R I E R P H D | 870 | Papio anubis |
| XP_025214301.1 | (+) | 1 | E L Q Q D D F Y A Y D E D G T R F S H D V T P I I L A A H C Q E Y E I V H T L L R K G A R I E R P H D | 931 | Theropithecus gelada |
| XP_011839769.1 | (+) | 1 | E L Q Q D D F Y A Y D E D G T R F S H D V T P I I L A A H C Q E Y E I V H T L L R K G A R I E R P H D | 931 | Mandrillus leucophaeus |
| XP_033060179.1 | (+) | 1 | E L Q Q D D F Y A Y D E D G T R F S H D V T P I I L A A H C Q E Y E I V H T L L R K G A R I E R P H D | 931 | Trachypithecus francoisi |
| XP_017744080.1 | (+) | 1 | E L Q Q D D F Y A Y D E D G T R F S H D V T P I I L A A H C Q E Y E I V H T L L R K G A R I E R P H D | 826 | Rhinopithecus bieti |
| XP_010357185.1 | (+) | 1 | E L Q Q D D F Y A Y D E D G T R F S H D V T P I I L A A H C Q E Y E I V H T L L R K G A R I E R P H D | 931 | Rhinopithecus roxellana |
| XP_023063874.2 | (+) | 1 | E L Q Q D D F Y A Y D E D G T R F S H D V T P I I L A A H C Q E Y E I V H T L L R K G A R I E R P H D | 931 | Piliocolobus tephrosceles |
| XP_018891488.1 | (+) | 1 | E L Q Q D D F Y A Y D E D G T R F S H D V T P I I L A A H C Q E Y E I V H T L L R K G A R I E R P H D | 931 | Gorilla gorilla gorilla |
| XP_003828441.1 | (+) | 1 | E L Q Q D D F Y A Y D E D G T R F S H D V T P I I L A A H C Q E Y E I V H T L L R K G A R I E R P H D | 931 | Pan paniscus |
| XP_024111625.1 | (+) | 1 | E L Q Q D D F Y A Y D E D G T R F S H D V T P I I L A A H C Q E Y E I V H T L L R K G A R I E R P H D | 931 | Pongo abelii |
| XP_030685496.1 | (+) | 1 | E L Q Q D D F Y A Y D E D G T R F S H D V T P I I L A A H C Q E Y E I V H T L L R K G A R I E R P H D | 931 | Nomascus leucogenys |
| XP_032023115.1 | (+) | 1 | E L Q Q D D F Y A Y D E D G T R F S H D V T P I I L A A H C Q E Y E I V H T L L R K G A R I E R P H D | 931 | Hylobates moloch |
| XP_039327066.1 | (+) | 1 | E L Q Q D D F Y A Y D E D G T R F S H D V T P I I L A A H C Q E Y E I V H T L L R K G A R I E R P H D | 931 | Saimiri boliviensis bolivi... |
| XP_032151593.1 | (+) | 1 | E L Q Q D D F Y A Y D E D G T R F S H D V T P I I L A A H C Q E Y E I V H T L L R K G A R I E R P H D | 931 | Sapajus apella |
| XP_017401411.1 | (+) | 1 | E L Q Q D D F Y A Y D E D G T R F S H D V T P I I L A A H C Q E Y E I V H T L L R K G A R I E R P H D | 931 | Cebus imitator |
| XP_021527350.1 | (+) | 1 | E L Q Q D D F Y A Y D E D G T R F S H D V T P I I L A A H C Q E Y E I V H T L L R K G A R I E R P H D | 931 | Aotus nancymaae |
| XP_008069472.1 | (+) | 1 | E L Q Q D D F Y A Y D E D G T R F S H D V T P I I L A A H C Q E Y E I V H T L L R K G A R I E R P H D | 845 | Carlito syrichta |
| XP_012514618.1 | (+) | 1 | E L Q Q D D F Y A Y D E D G T R F S H D V T P I I L A A H C Q E Y E I V H T L L R K G A R I E R P H D | 931 | Propithecus coquereli |
| XP_020142255.1 | (+) | 1 | E L Q Q D D F Y A Y D E D G T R F S H D V T P I I L A A H C Q E Y E I V H T L L R K G A R I E R P H D | 931 | Microcebus murinus |
| XP_003780834.1 | (+) | 1 | E L Q Q D D F Y A Y D E D G T R F S H D V T P I I L A A H C Q E Y E I V H T L L R K G A R I E R P H D | 931 | Otolemur garnettii |
| XP_007936274.1 | (+) | 1 | E L Q H D D F Y A Y D E D G T R F S H D V T P I I L A A H C Q E Y E I V H T L L R K G A R I E R P H D | 929 | Orycteropus afer afer |
| XP_006887384.1 | (+) | 1 | E L Q Q D D F Y A Y D E D G T R F S H D V T P I I L A A H C Q E Y E I V H T L L R K G A R I E R P H D | 929 | Elephantulus edwardii |
| XP_004709071.1 | (+) | 1 | E L Q Q D D F Y A Y D E D G T R F S H D V T P I I L A A H C Q E Y E I V H T L L R K G A R I E R P H D | 729 | Echinops telfairi |
| XP_003415645.1 | (+) | 1 | E L Q Q D D F Y A Y D E D G T R F S H D V T P I I L A A H C Q E Y E I V H T L L R K G A R I E R P H D | 929 | Loxodonta africana |
| XP_037697124.1 | (+) | 1 | E L Q Q D D F Y A Y D E D G T R F S H D V T P I I L A A H C Q E Y E I V H T L L R K G A R I E R P H D | 905 | Choloepus didactylus |
| XP_028903564.1 | (+) | 1 | E I Q L D D F Y A Y D E D G T R F S H D V T P I I L A A H C Q E Y E I V H T L L R K G A R I E R P H D | 972 | Ornithorhynchus anatinus |
| XP_038617812.1 | (+) | 1 | E I Q L D D F Y A Y D E D G T R F S H D V T P I I L A A H C Q E Y E I V H T L L R K G A R I E R P H D | 912 | Tachyglossus aculeatus |
| XP_022368885.1 | (+) | 1 | E L Q Q D D F Y A Y D E D G T R F S H D V T P I I L A A H C Q E Y E I V H T L L R K G A R I E R P H D | 932 | Enhydra lutris kenyoni |
| XP_004427467.1 | (+) | 1 | E L Q Q D D F Y A Y D E D G T R F S H D V T P I I L A A H C Q E Y E I V H T L L R K G A R I E R P H D | 932 | Ceratotherium simum sim... |

Supplemental Figure 2: Amino acid protein sequence alignment of three major homologous TRPC channels, TRPC3, TRPC6, and TRPC7. The primary regulating threonine (T221 – TRPC6; T150 – TRPC3, and T166 – TRPC7 are conserved and surrounded by highly conserved sequences.

```
sp|Q9Y210|TRPC6_HUMAN      DNRLAHRRQTVLREKGRRLANRGPAYMFSDRSTSLSIEEERFLDAAEYGNIPVVRKMLEE 120
sp|Q13507|TRPC3_HUMAN      -----------MREKGRRQAVRGPAFMFNDRGTSLTAEEERFLDAAEYGNIPVVRKMLEE 49
sp|Q9HCX4|TRPC7_HUMAN      TFKNMQRRHTTLREKGRRQAIRGPAYMFNEKGTSLTPEEERFLDSAEYGNIPVVRKMLEE 65
                                      :****** * ****:**.::.***: *******:***************

sp|Q9Y210|TRPC6_HUMAN      CHSLNVNCVDYMGQNALQLAVANEHLEITELLLKKENLSRVGDALLLAISKGYVRIVEAI 180
sp|Q13507|TRPC3_HUMAN      SKTLNVNCVDYMGQNALQLAVGNEHLEVTELLLKKENLARIGDALLLAISKGYVRIVEAI 109
sp|Q9HCX4|TRPC7_HUMAN      SKTLNFNCVDYMGQNALQLAVGNEHLEVTELLLKKENLARVGDALLLAISKGYVRIVEAI 125
                           .::**.***************.*****:**********:*:*******************

sp|Q9Y210|TRPC6_HUMAN      LSHPAFAEGKRLATSPSQSELQQDDFYAYDEDGTRFSHDVTPIILAAHCQEYEIVHTLLR 240
sp|Q13507|TRPC3_HUMAN      LNHPGFAASKRLTLSPCEQELQDDDFYAYDEDGTRFSPDITPIILAAHCQKYEVVHMLLM 169
sp|Q9HCX4|TRPC7_HUMAN      LNHPAFAQGQRLTLSPLEQELRDDDFYAYDEDGTRFSHDITPIILAAHCQEYEIVHILLL 185
                           *.**.** .:**: ** :.**::************** *:**********:**:** **

sp|Q9Y210|TRPC6_HUMAN      KGARIERPHDYFCKCNDCNQKQKHDSFSHSRSRINAYKGLASPAYLSLSSEDPVMTALEL 300
sp|Q13507|TRPC3_HUMAN      KGARIERPHDYFCKCGDCMEKQRHDSFSHSRSRINAYKGLASPAYLSLSSEDPVLTALEL 229
sp|Q9HCX4|TRPC7_HUMAN      KGARIERPHDYFCKCNECTEKQRKDSFSHSRSRMNAYKGLASAAYLSLSSEDPVLTALEL 245
                           ***************.:* :**::*********:******** ***********:*****

sp|Q9Y210|TRPC6_HUMAN      SNELAVLANIEKEFKNDYKKLSMQCKDFVVGLLDLCRNTEEVEAILNGDVETLQ--SGDH 358
sp|Q13507|TRPC3_HUMAN      SNELAKLANIEKEFKNDYRKLSMQCKDFVVGVLDLCRDSEEVEAILNGDLESAEPLEVHR 289
sp|Q9HCX4|TRPC7_HUMAN      SNELARLANIETEFKNDYRKLSMQCKDFVVGVLDLCRDTEEVEAILNGDVNFQV--WSDH 303
                           ***** *****.******:************:*****::**********::      .:
```
